# The C-terminal tails of GroEL and its mitochondrial and chloroplastic homologs adopt polyproline II helices

**DOI:** 10.1101/2024.05.27.596059

**Authors:** Cristian Segura Rodríguez, Rubén López-Sánchez, Douglas Vinson Laurents

## Abstract

The chaperonin GroEL and its mitochondrial and chloroplastic homologs mHsp60 and Cpn60 are large barrel-like oligomeric proteins. Chaperonins facilitate folding by isolating nascent chains in their hollow interior and undergoing conformational transitions driven by ATP hydrolysis. Due to their vital importance, the structure of GroEL and its homologs have been extensively studied by X-ray crystallography and CryoEM, revealing one or two rings each of which contains seven subunits. Each subunit has three folded domains and a twenty-four residue C-terminal extension. Whereas this C-terminal tail has been reported to bind and stimulate the folding of the client protein, it appears to be invisible or blurry, which suggests disorder. The objective of this study is to characterize conformational preferences in the C-terminal tails of GroEL, mHsp60 and representative Cpn60s using circular dichroism and nuclear magnetic resonance spectroscopies. The tails of GroEL and mHsp60 consist of two segments. The first is rich in residues typical of intrinsically disordered proteins and the second segment consists exclusively (GroEL) or almost entirely (mHsp60) of Gly and Met residues. The spectroscopic results evince that these C-terminal extensions are not wholly disordered but adopt polyproline II helices whose populations are higher in the second Gly/Met-rich segment. Whereas the C-terminal segments of chloroplastic chaperonins are Gly-poor, they are rich in proline and also adopt polyproline II helix conformations. These results provide insight into the function of chaperonin C-terminal tails.

## Introduction

Chaperonins are protein folding machines found in all kingdoms of life^1,2^. Eubacteria chaperonins, as exemplified by the well characterized protein GroEL from *E. coli* consist of seven subunits, each containing three folded domains. These oligomerize to form a homo-heptamer with a hollow interior. Unfolded client proteins enter the barrel and bind to its interior surface. The open barrel is then capped by GroES, a smaller homoheptameric protein that acts like a lid. Conformational changes in the complex, which are influenced by ATP hydrolysis, GroES binding and the dimerization of the GroEL_7_·GroES_7_ complex, aid the proper folding of client proteins^2^. GroEL homologs are present in mitochondria (called Hsp60, sequence identity versus GroEL ∼60%) with a similar structure^3^, and chloroplasts (called Cpn60, sequence identity to GroEL ∼ 60%)^4^. In addition to the ordered domains, both GroEL and Hsp60 contain disordered C-terminal tails of about 25 residues in length which appear invisible or diffuse by X-ray crystallography and CyroEM and are predicted to be disordered by AlphaFold (**Figure 1**). Nevertheless, these tails contribute to substrate folding^5^ and form a barrier that blocks substrates from escaping through the bottom of the GroEL barrel^6^. In GroEL and Hsp60, these tails consist of several charged, short or polar residues followed by a ∼19 residue segment with a remarkably high glycine and methionine content (**Sup. Table 1**). Recently, a structural class of proteins containing glycine-rich polyproline II helical bundle domains has emerged^7^. Moreover, the C-terminal Gly/Met-rich segments of GroEl and mHsp60 resemble Abductin, an elastic protein present in the hinge ligament of bivalves^8^, which adopts polyproline II (PPII) helix conformations^9^. This led us to wonder whether the C-terminal tails of GroEL and mHsp60 might also adopt PPII helical conformations, and here we will test this hypothesis.

**Figure 1.**
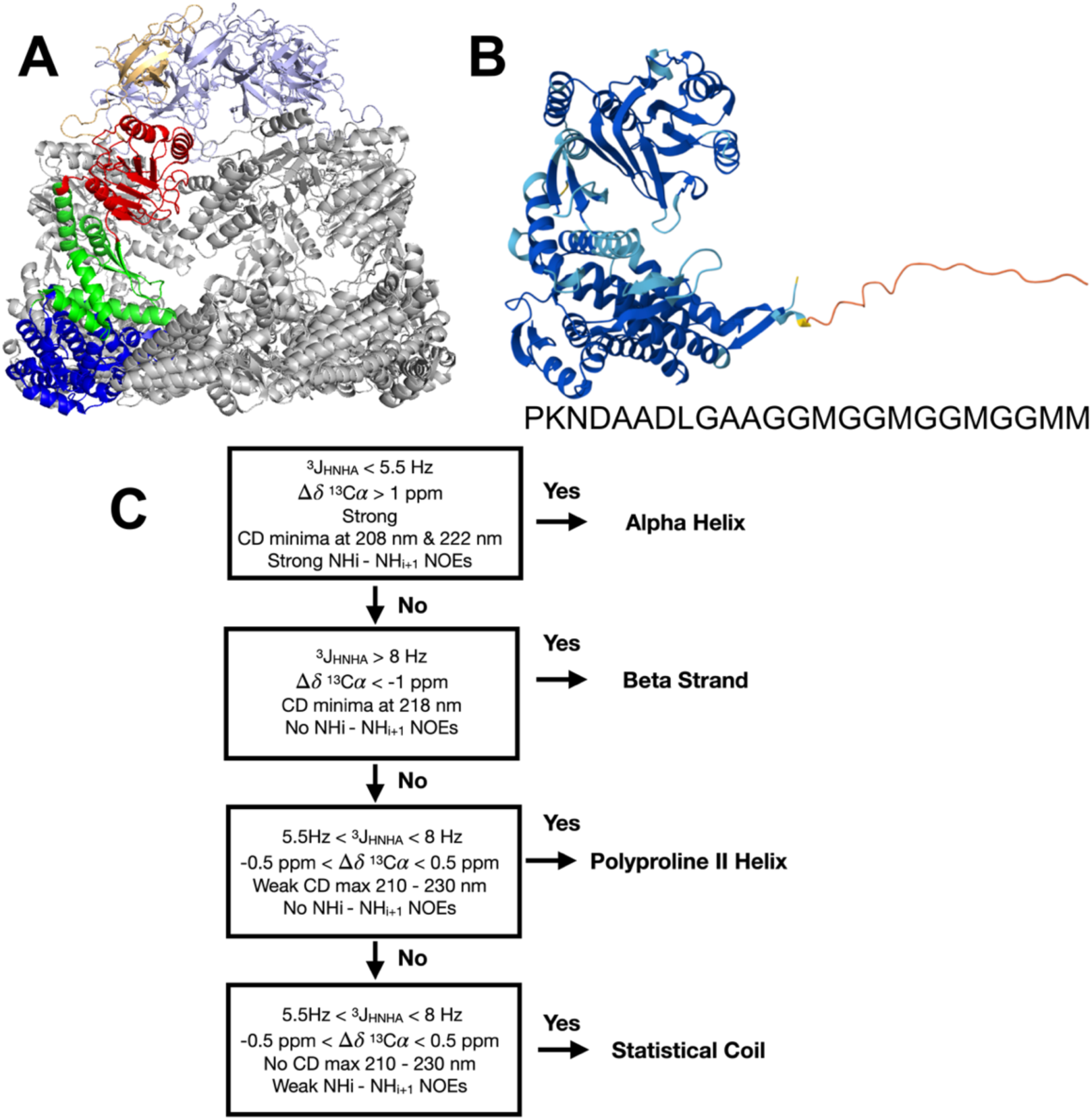
The “disordered” C-terminal segment in GroEL/mHsp60 and spectral criteria for its conformational characterization. **A**. Structure of mHsp60/mHsp10 complex: PDB=6MRD (ref. #3). Six mHsp10 subunits are tinted blue/white, the seventh is colored gold. Six mHsp60 subunits are colored gray, the seventh is colored red, green and blue for its apical, intermediate and equatorial domains, respectively. **B**. AlphaFold2 structural model (https://alphafold.ebi.ac.uk/entry/P0A6F5) of *E. coli* GroEL (one subunit). Residues whose conformation is predicted with high confidence are colored blue (the three folded domains) and those with low/very low confidence are colored orange and red; namely the C-terminal segment, whose amino acid sequence is written. **C**. Criteria from NMR data and CD spectral features which can distinguish α-helices, β-strands, polyproline II helices and statistical coil ensembles. These criteria are based on those of Ma *et al.*^10^ but substituting the ^1^HαΔδ for ^13^CαΔδ, as the latter are more reliable^11^.

The chloroplastic chaperonins, which are key for RuBisCO folding^12^, contain two types of subunits, called α and β, have been reported to form heptamers^13^; which then combine to form heterodecatetramers^14^. Like the distinct subunits of cytoplasmatic chaperonins^4^, it is possible that the α and β subunits of Cpn60 underwent divergent evolution to optimize function^15^. The chloroplastic subunits also have C-terminal extensions predicted to be disordered by AlphaFold, but unlike the tails of GroEL and mHsp60, they are glycine-poor but are often rich in proline residues (**Sup. Table 1**). Characterizing these segments is a second goal of this study.

## Results

### The Gly-rich stretch of GroEL and mHsp60 contains a high percentage of PPII helix

To test for the presence of polyproline II helix, we studied short peptides corresponding to just the Gly/Met-rich stretch right at the C-terminus of GroEL and mHsp60. The NMR spectra of these peptides, called GroELCtS and mHsp60CtS, show poor ^1^HN and ^13^Cα chemical shift dispersion and degenerate ^1^Hα glycine signals (**Sup. Fig. 1**). Moreover the ^13^Cα conformational chemical shifts are very small. These chemical shift data indicate an absence of α-helix and β-strand conformations (**Figure 2A,B**). The measured ^3^J_HNHα_ couplings are consistent with extended conformations.

**Figure 2.**
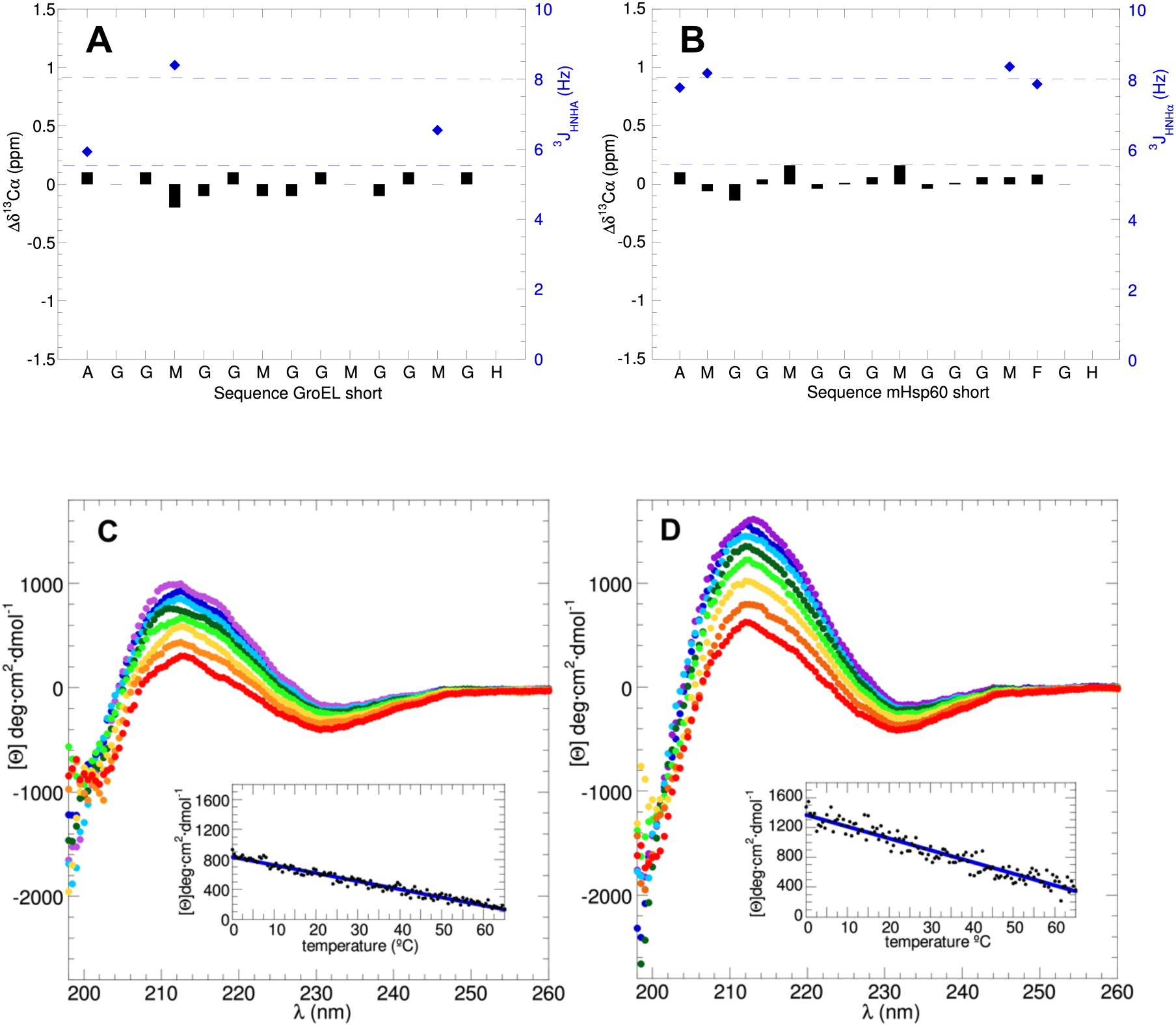
NMR spectral parameters and CD spectra of *GroEL and mHsp60* C-terminal segments. **A**. GroELCtS ^13^Cα conformation chemical shifts (black bars, left y-axis) and ^3^J_HNHα_ (blue diamonds, right y-axis). As in (**B**), the latter are measured for non-overlapping, non-Glycine residues. **B**. mHsp60CtS ^13^Cα conformation chemical shifts (black bars, left y-axis) and ^3^J_HNHα_ (blue diamonds, right y-axis). **C**. Far UV-CD spectra of GroELCtS recorded at 0 °C (violet), 5°C (blue), 10°C (celeste), 17°C (dark green), 25°C (light green), 37°C (yellow), 50°C (orange), 65°C (red). *Inset:* Terminal denaturation of the GroELCtS. The fit of a linear equation (blue line) to the data (black points) is [0]_217nm_= 835 – 10.8 (T °C) R= 0.98. **D**. Far UV-CD spectra of mHsp60CtS recorded at 0 °C (violet), 5°C (blue), 10°C (celeste), 17°C (dark green), 25°C (light green), 37°C (yellow), 50°C (orange), 65°C (red). *Inset:* Heat denaturation of the C-terminal segment of mHsp60CtS. The fit of a linear equation (blue line) to the data (black points) is [0]_217nm_= 13200 – 15.8(T °C) R= 0.96.

The far UV-CD spectra of GroELCtS and mHsp60CtS, recorded over a range of temperatures from 0 to 65°C, are shown in **Fig. 2C,D**. At low temperatures, the spectra show a maximum near 212 nm which is characteristic of polyamide PPII helices^16,17^. Upon heating, this maximum weakens in intensity. The CD spectral bands are stronger for Hsp60CtS than GroELCtS. This could be attributed to a greater number of glycine residues in the former, as the intrinsic PPII forming propensity of Gly is higher than Met^18,19^. Based on their maximum CD signals of 1600 deg·cm^2^·dmol^-1^ and 1000 deg·cm^2^·dmol^-1^, the populations of PPII helix at 0 °C are estimated to be 47% and 43% for Hsp60CtS and GroELCtS, respectively (see **Experimental Procedures**). At 37°C, the physiological relevant temperature, the %PPII helix would be 43% and 41% for Hsp60CtS and GroELCtS, respectively.

Next, the signal at 217 nm was recorded as the sample was heated at 1 °C per minute up to 65 °C. Instead of the sigmoidal transition typically observed for folded protein, GroELCtS and mHsp60CtS show a linear loss of signal upon heating (**Fig. 2C,D insets**). This is consistent with a non-cooperative PPII helix to statistical coil transition, which has been previously observed for other isolated PPII helices^20^ or partly populated PPII helices in intrinsically disordered proteins^21^.

### Complete C-terminal tails contain smaller populations of polyproline II helix

To further corroborate and assess the position of the PPII helix, we also characterized peptides, called GroELCtC and mHsp60CtC, corresponding to the complete segments of GroEL and mHsp60 that are invisible to X-ray crystallographic and CryoEM analyses. The assigned NMR spectra of GroELCtC and mHsp60CtC shown respectively in **Sup. Figures 2 & 3** reveal a lack of ^1^HN and ^13^Cα chemical shift dispersion. This is characteristic of an absence of α-helical and β-strand structure^11,22^. Whereas two different ^1^Hα signals were seen for H-bonded glycine residues in a folded protein consisting of six PPII helices^23^, the two glycine ^1^Hα signals are degenerate in all GroELCt and mHsp60Ct peptides (**Sup. Figures 1, 2 & 3**). This is relevant because the seven C-terminal segments extending from the equatorial domains towards the barrel interior might associate to form a structured bundle of polyproline II helices. The lack distinct glycine ^1^Hα signals suggests they do not. The small ^13^Cα conformational chemical shifts and ^3^J_HNHα_ values in the range of 5.5 to 8.0 Hz^24,20,25^ rule out the presence of α-helices or β-strands (**Sup. Figure 4 A&B**). The presence of weak maxima in the CD spectra of GroELCtC and mHsp60CtC (**Sup Fig. 4 C&D**) indicate the presence of polyproline II helix. These peaks become weaker upon heating manifesting a partial PPII helix to coil transition which has been observed previously for other PPII helices^26,27^. The difference spectra, which were calculated by subtracting spectra recorded at two different temperatures (**Sup. Fig 4 C&D insets**), are very similar to the CD spectra of peptides^28,29^, collagen^30^ and proteins^17^ known to adopt PPII helices.

### Chloroplast homologs of GroEL, which are Proline-rich, also adopt PPII conformations

The NMR spectra of *A. thaliana* and wheat Cpn60α C-terminal segments, shown in **Sup. Fig. 5**, reveal a poor ^1^HN chemical shift dispersion which is consistent with a lack of well ordered α-helical or β-sheet structures. There are no detectable ^1^HN_i_-^1^HN_i+1_ or ^1^HN_i_-^1^HN_i+3_ NOE signals which would be characteristic of α-helices. The ^3^J_HNHα_ coupling constants and small or slightly negative ^13^Cα conformational chemical shifts are indicative of statistical coil and extended (β-strand or PPII helical) conformations (**Figure 3A,B**). Similar results are seen for the C-terminal segments of *Arabadopsis thaliana* and wheat Cpn60β subunits (**Sup. Figure 6A**). The CD spectra of peptides corresponding to the C-terminal segments of *Arabadopsis thaliana* and wheat Cpn60α recorded at 5, 25, 37 and 65 °C are shown in **Figure 3C,D**. These spectra show a weak maximum near 224 nm at 5°C which is suggestive of PPII helical conformations. Except for minor differences in the position of the peak maxima, which could be ascribed to a difference in the content of proline and aromatic residues, the results for the C-terminal segments of *Arabadopsis thaliana* and wheat Cpn60β subunits are similar (**Sup. Figure 6B,C**). Upon heating, the spectra show significant changes including a decreased signal around 220 nm. Like the difference spectra of GroELCtC and mHsp60CtC, the chloroplastic difference spectra closely resemble typical PPII helical spectra. Based on the CD data, the populations of PPII helix in the C-terminal segments of *A. thaliana* and wheat Cpn60α are about 30% and are slightly higher (32% and 34%) for *A. thaliana* and wheat Cpn60β, respectively, at 25°C.

**Figure 3.**
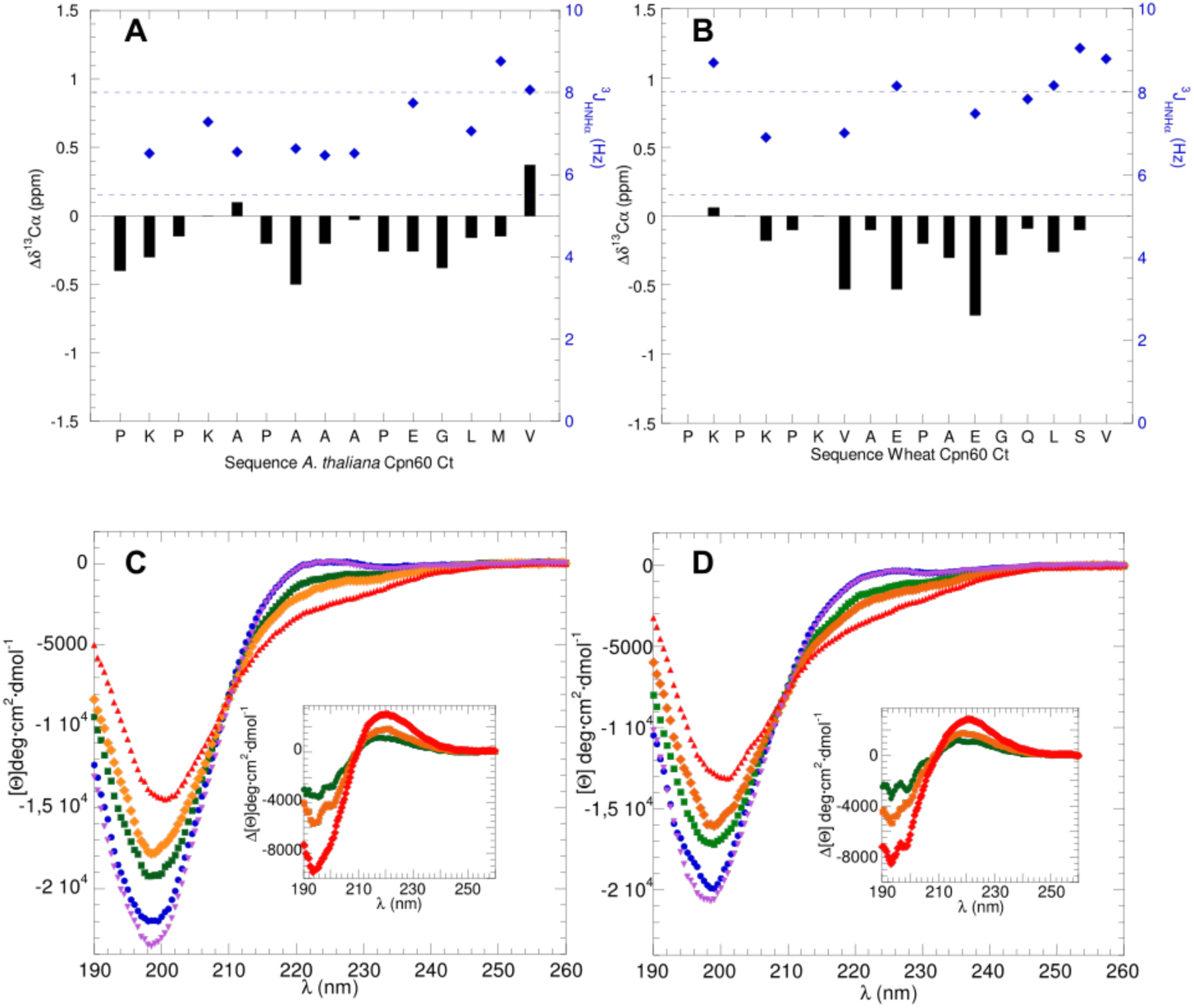
NMR spectral parameters and CD spectra of *Arabadopsis thaliana* and Wheat Cpn60 C-terminal segments. ^13^Cα conformational chemistry shifts (black bars, left y-axis) and ^3^J_HNHα_ coupling constants (blue diamonds, right y-axis) for the *A. thaliana* (panel **A**) and wheat (panel **B**) Cpn60 C-terminal segments. Blue dashed horizontal lines mark the range of constants (between 5.5 & 8.0 Hz) expected for statistical coil or polyproline II conformations. Far UV-CD spectra recorded at 5°C(blue), 25°C(green) 37°C(orange) and 65°C(red) and after recooling to 5°C (purple) for the *A. thaliana* (panel **C**) and wheat (panel **D**) Cpn60α C-terminal segments. Insets show the difference CD spectra for 5°C-25°C (green), 5°C-37°C(orange) and 5°C-65°C(red).

## Discussion

The main finding of this study is that peptides corresponding to the C-terminal extensions of eubacterial, mitochondrial and chloroplastic chaperonins, which appear disordered in cryoEM and X-ray crystallographic analyses, adopt high populations of polyproline II helix in aqueous solution. This is consistent with the high intrinsic propensity of glycine residues to adopt isolated PPII helices^31^ as well as PPII helical bundles in synthetic peptides^32^, natural protein domains^33–35^ and quasi infinite honeycomb networks^36^.

Despite the sequence similarity to the “snow flea” antifreeze protein^23^ and abductin^9^, which contain glycine-rich polyproline II helices assembled into bundles, the C-terminal chaperonin peptides appear adopt isolated polyproline II helices under the conditions employed here. Nevertheless, since the seven C-terminal tails would be present at concentrations of dozens of mM and subject to crowding effects^37^ within chaperonin barrel, the possibility that they may associate under physiological conditions can not be ruled out. The associations between PPII helices of the C-terminal tails could plug the bottom of the chaperonin barrel and prevent client proteins from escaping^6^. Incidentally, the association of the PPII helices formed by Gly-rich C-terminal segments of GroEL into putative PPII helical bundle structures could explain why they are much more resistant to protease K degradation than the C-terminal segments of chloroplastic Cpn60^38^. The latter, being Pro-rich but Gly-poor, can form isolated PPII helices but cannot associate^7^.

The results show significant populations of PPII helix in both the α and β subunits of Cpn60. Sequence comparison of Cpn60 α and β subunits (**Sup. Table 1**) suggests that the C-terminal extensions of their α subunits often start with three cationic residues and end with two hydrophobic residues. By contrast, β-subunits frequently begin with two anionic residues and end in a SGYGY motif that is reminiscent of the GY segments of TDP-43 and FUS proteins, which is known to promote π/π and cation/π interactions^39,40^. These differences suggest that the C-terminal tails of the α and β subunits may have specialized through evolution to fold client proteins rich in anionic and aliphatic (Cpn60α subunit) or cationic and aromatic (Cpn60β subunit).

## Experimental Procedures

### Peptides

Peptides corresponding the complete GroEL and mHsp60 C-terminal segments with sequences: acPKNDAADLGAAGGMGGMGGMGGMMKKK and acPKEEKDPGMGAMGGMGGGMGGGMFKKK, respectively, were obtained from CASLO ApS, Denmark. These “<u>c</u>omplete” C-terminal peptides, are abbreviated as GroELCtC and mHsp60CtC, respectively. These peptides contain three extra Lys residues at the C-terminus to promote solubility. All peptides were N-acetylated as indicated by “ac” to mimic the lack of charge in the context of the full length protein.

Shorter peptides with just the final Gly/Met rich stretch of these peptides were also obtained from Calso ApS. The peptides, named GroELCtS and mHsp60CtS (the last “S” standing for short) have the sequences: acAGGMGGMGGMGGMMH and acAMGGMGGGMGGGMFGH, respectively. For these peptides, one C-terminal His residue was added to promote solubility.

The N-acetylated peptides acPKPKAPAAAPEGLMV, acPKPKPKVAEPAEGQLSV, acEPEPVPVGNPMDNSGYGY and acEPEAAPLANPMDNSGFGY which correspond to the C-terminal segments of *Arabadopsis thaliana* Cpn60α, wheat Cpn60α, *Arabadopsis thaliana* Cpn60β, and wheat Cpn60β subunits respectively, were purchased from the National Center for Biotechnology (CNB, CSIC) Cantoblanco, Spain.

The sequence and purity of all peptides was verified by mass spectrometry and NMR spectroscopy. Sample concentrations were measured by integration of 1D ^1^H peptide NMR signals to that of a known concentration of sodium trimethylsilylpropanesulfonate, which was also used as the internal chemical shift standard.

### NMR Spectroscopy

A series of NMR spectra; namely: 2D ^1^H-^1^H COSY, TOCSY and NOESY and 2D ^1^H-^13^C HSQC and ^1^H-^15^N HSQC were recorded on 1 – 3 mM peptide samples in 10 mM K_2_HPO_4_ buffer (pH 6) and 5°C on the 600 MHz (^1^H) Bruker AvanceNeo spectrometer belonging to the “Manuel Rico” NMR laboratory, IQF/CSIC. The spectral parameters are summarized in **Sup. Table 2**. The spectra were assigned and analyzed to obtain ^13^Cα chemical shifts and ^3^J_ΗΝHα_ coupling constants. The ^13^Cα chemical shifts reveal conformational tendencies^41^ when compared to reference ^13^Cα chemical shifts predicted for a structureless peptide, which were obtained for each peptide and set of experimental conditions using previously reported parameters^42^ (**Figure 1**). ^3^J_HNHα_ couplings also discriminate between α-helical, β-strand, and statistical coil or polyproline II conformations^43^ (**Figure 1**).

### Circular Dichroism Spectroscopy

Far UV circular dichroism (CD) spectra were recorded on a Jasco 810 spectropolarimeter equipped with a Peltier temperature control unit at 190 – 260 nm or 198 – 260 nm with a 50 nm/min scan speed, 1.67 nm bandwidth, peptide concentrations ranging from 130 to 300 µM in 10 mM K_2_HPO_4_ buffer prepared in milliQ water at pH 6. Ten scans were recorded and averaged per spectrum. A reference buffer spectrum was subtracted.

Thermal unfolding experiments were monitored by CD at 217 nm from 0°C to 65°C employing a 1°C/min heating rate and a 1.0 nm bandwidth. Spectra were recorded at 0°C before and after the thermal denaturation experiment to test reversibility. The empirical equation of Stellwagen and co-workers^44^:

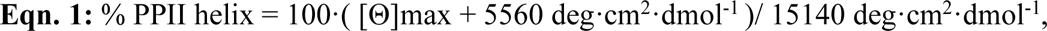

where [Θ]max is the maximum in the spectrum between 210 and 230 nm in deg·cm^2^·dmol^-1^, was used to estimate the population of PPII helix.

## Supporting Information Available

Tables with sequence information, NMR spectral parameters, Figures with NMR & CD spectra and spectral results are included in the Supporting Information.

## Acknowledgements

We are grateful to Dr. I. Tascón, Prof. I. Ubarretxena, and Prof. J. M. Valpuesta for helpful conversation and encouragement.

## Funding and Additional Information

This study is part of projects PID2019 109306RB-I00 and PID2022-137806OB-I00, funded by the Spanish Ministry of Science, Innovation and Universities: MICIN/AEI/10.13039/501100011033/FEDER, UE. NMR experiments were performed in the “Manuel Rico” NMR laboratory (LMR) of the Spanish National Research Council (CSIC), a node of the Spanish Large-Scale National Facility (ICTS R-LRB).

## Table of Contents Graphic

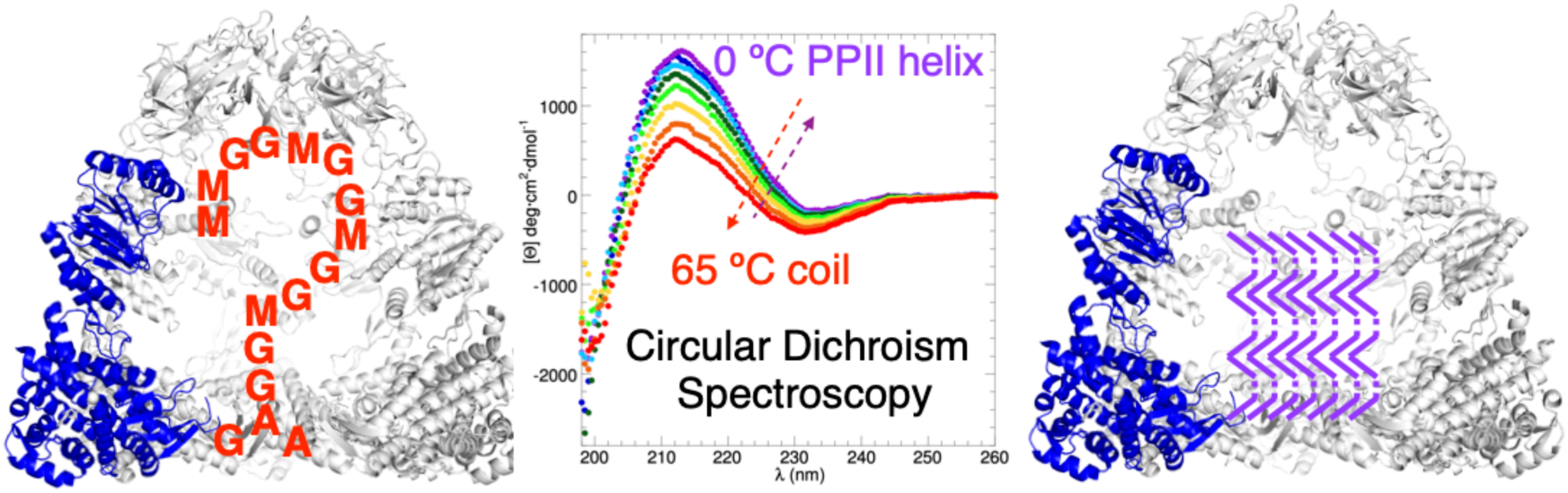

## Supporting Information

**Supporting Table 1.**
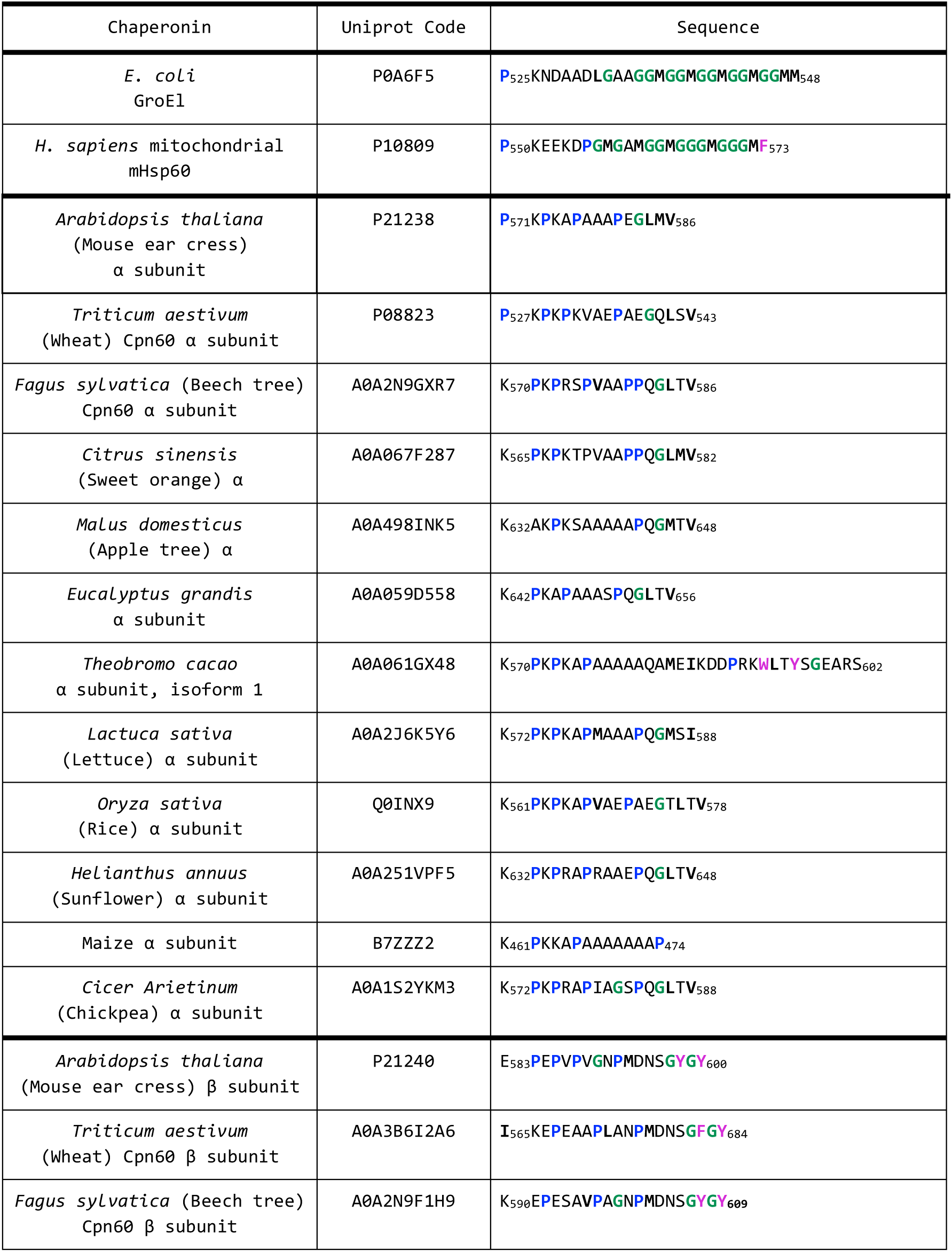
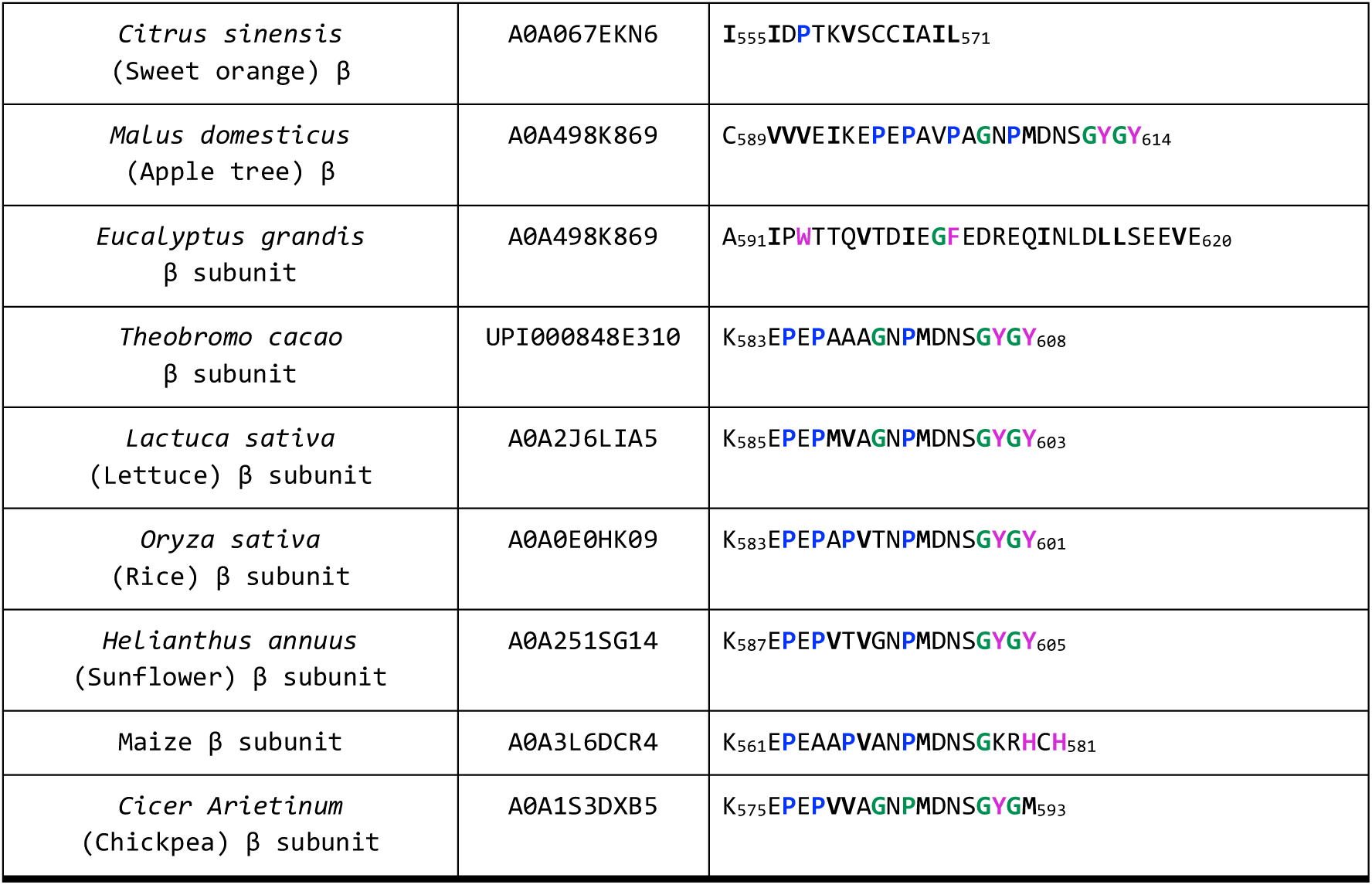
Aminoacid sequences of C-terminal disordered segments in *E. coli* GroEL, human mitochondrial mHsp60 and representative chloroplastic Cpn60 α and β subunits. Glycine, proline, aliphatic and aromatic residues are colored green, blue, bold black and purple, respectively.

**Supporting Table 2:** NMR Spectral Parameters.

| Type of Spectrum | Number of Scans | Sweep Width (ppm) | Matrix Size | Mixing Time (ms) |
| --- | --- | --- | --- | --- |
| 1D $^1\text{H}$ | 64 | 12 | - | - |
| 2D $^1\text{H}$, $^1\text{H}$ COSY | 16 | 11 x 11 | 8192 x 336 | - |
| 2D $^1\text{H}$, $^1\text{H}$ TOCSY | 16 | 11 x 11 | 2048 x 256 | 40, 60 or 100* |
| 2D $^1\text{H}$, $^1\text{H}$ NOESY | 40 | 11 x 11 | 2048 x 288 | 100-120 |
| 2D $^1\text{H}$-$^{13}\text{C}$ HSQC | 144 | 10.4 x 50 (center 40 ppm) | 1024 x 80 | - |
| 2D $^1\text{H}$-$^{15}\text{N}$ HSQC† | 336 | 10 x 23 | 2048 x 128 | - |

\* The default TOCSY mixing time was 60 ms, but some additional experiments were run using shorter or longer mixing times to distinguish and assign longer side chains.
† Recorded only for the longer peptides, namely: GroELCtC and mHsp60CtC.

**Supporting Figure 1:**
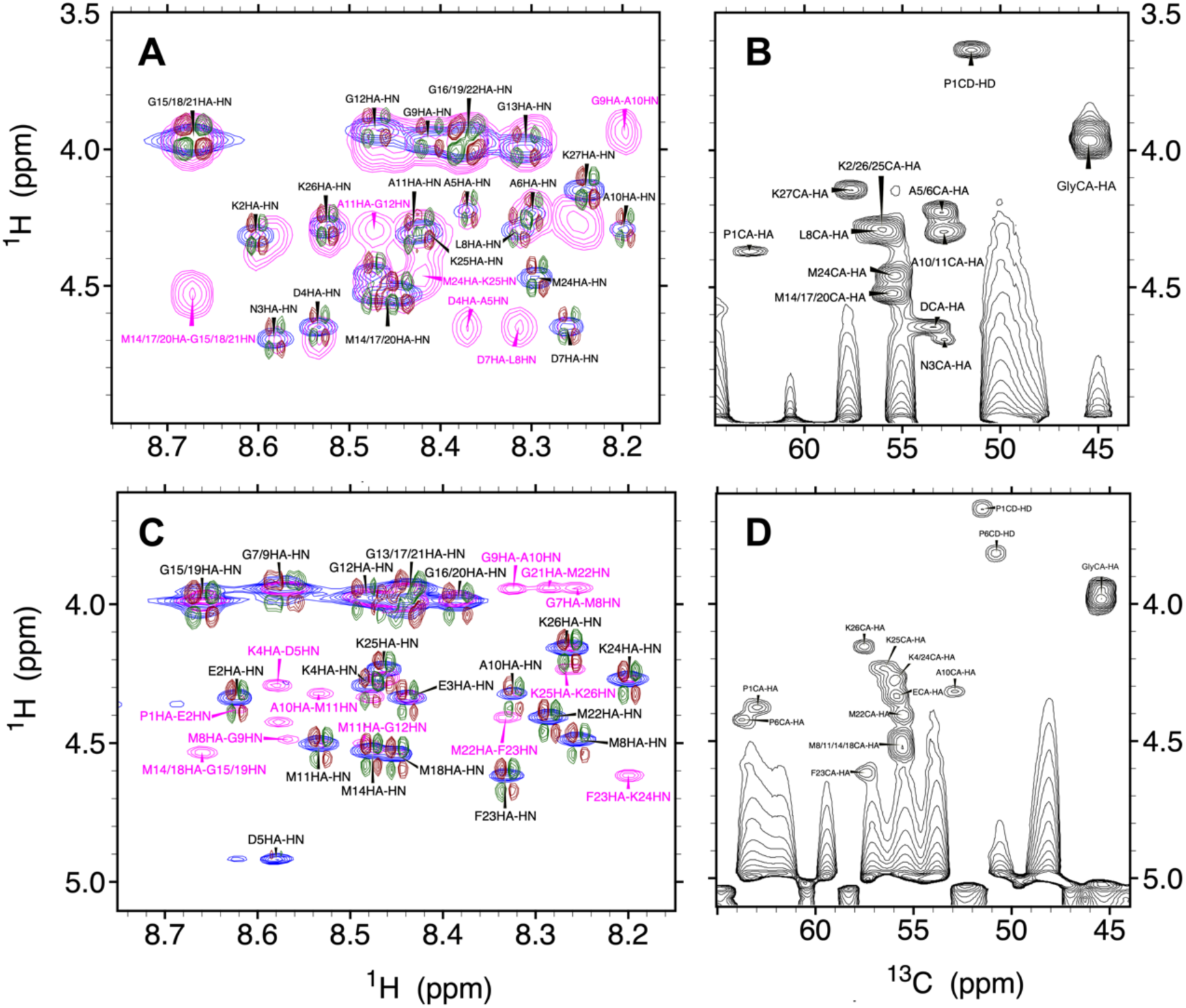
NMR spectra of GroELCtS and mHsp60CtS. **A.** GroELCtC 2D ^1^H-^1^H COSY (maroon/green), TOCSY (blue) and NOESY (magenta) NMR signals of the HN/Hα region are shown. Intra-residual crosspeaks are labeled in black and inter-residual signals are labeled in magenta. **B.** 2D ^1^H-^13^C HSQC signals from the ^13^Cα/^1^Hα region. Overlapped glycine signals are labeled “Gly” and overlapped Asp are labeled with a “D”. **C.** mHsp60 CtC 2D ^1^H-^1^H COSY (maroon/green), TOCSY (blue) and NOESY (magenta) signals of the HN/Hα region are shown. Intra-residual crosspeaks are labeled in black and inter-residual signals are labeled in magenta. **D.** mHsp60 CtC 2D ^1^H-^13^C HSQC signals from the ^13^Cα/^1^Hα region. Overlapped glycine signals are labeled “Gly”.

**Supporting Figure 2:**
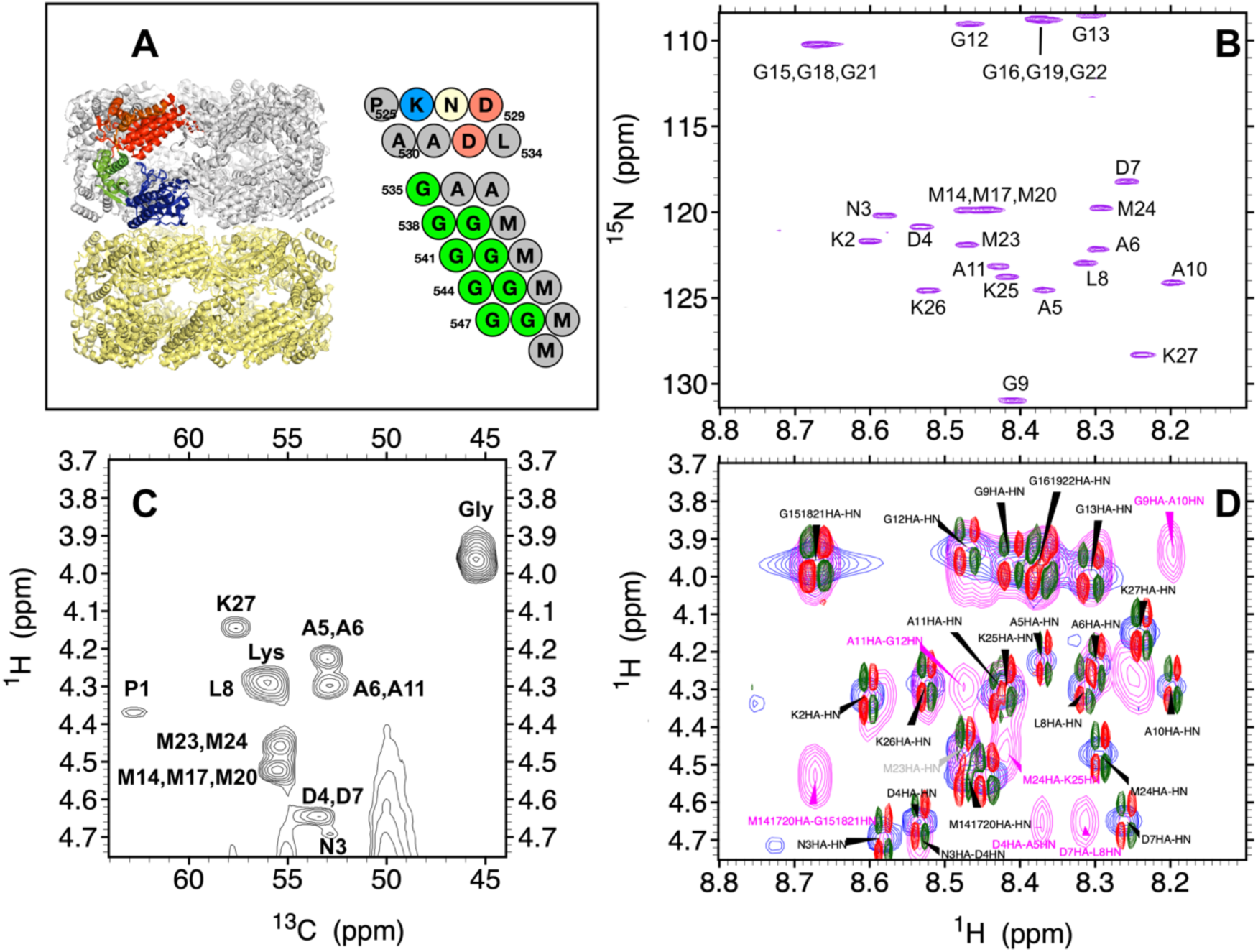
NMR spectra of GroEL C-terminal Segment. **A.** (*left*) Structure of GroEL (PDB 2EU1) showing seven subunits of the lower barrel in gold and six subunits of the upper barrel in silver. The apical, intermediate and equatorial domains of one subunit are shown in red, green and blue, respectively. (*right*) The residues of the GroEL C-terminal tail, which are invisible to CryoEM or X-ray diffraction. **B.** ^1^H-^15^N HSQC spectrum of the C-terminal tail of GroEL. The signal of G9 is folded. **C.** ^1^HN-^1^Hα region of 2D ^1^H-^1^H COSY (maroon/dark green), TOCSY (blue) and NOESY (magenta) spectra. Inter-residue crosspeaks are colored maroon and intra-residues crosspeaks are colored black. **D.** ^1^HN-^1^Hα region of 2D ^1^H-^13^C HSQC spectrum. Peaks with overlapped signals from multiple residues are labeled with the three-letter code (*e.g.* Gly, Lys).

**Supporting Figure 3:**
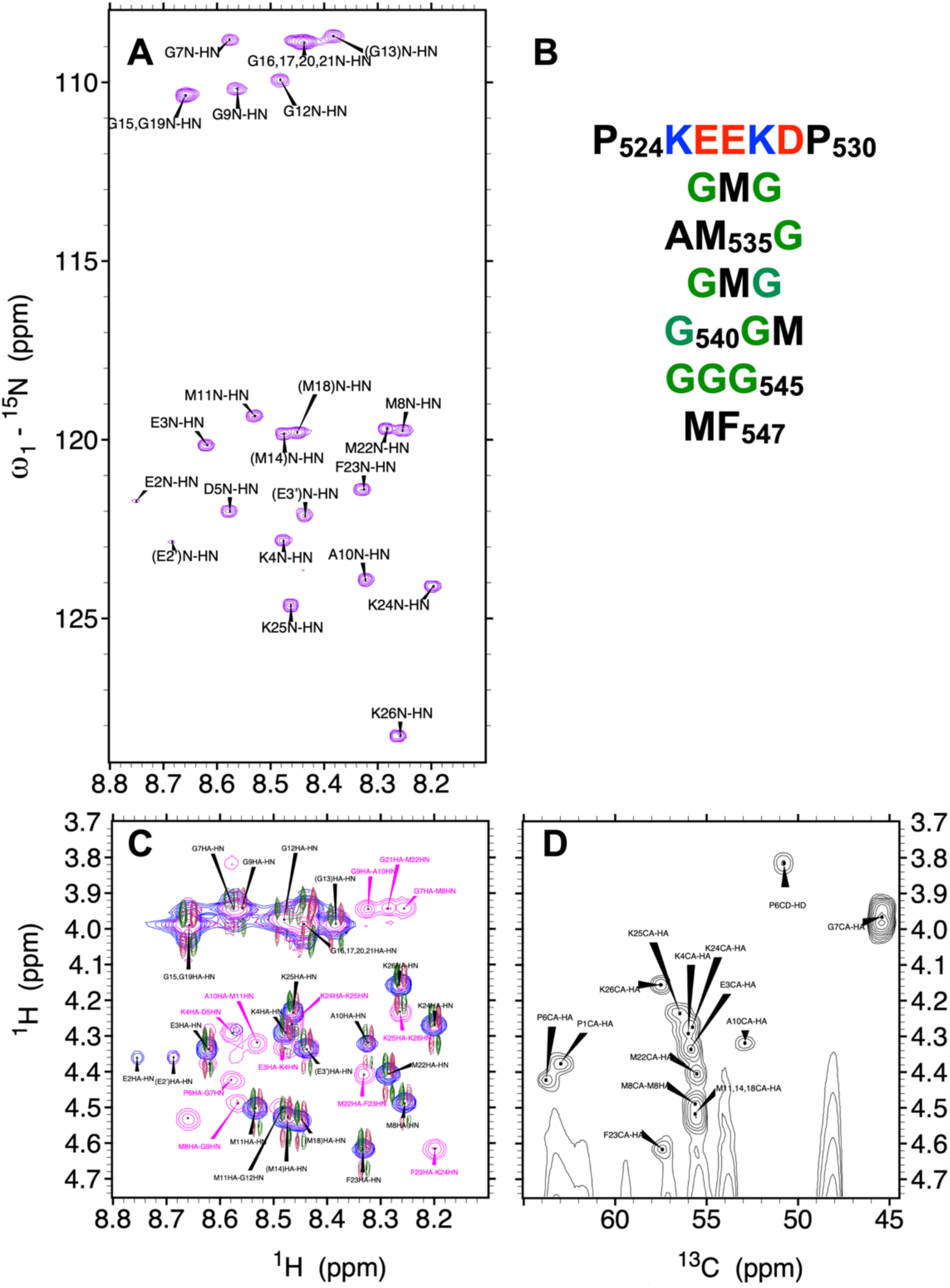
NMR spectra of mHsp60 C-terminal Segment. **A.** ^1^H-^15^N HSQC spectrum of the C-terminal tail of mHsp60. Ambiguous assignments are shown in parentheses. **B.** The residues of the mHsp60 C-terminal tail, which are invisible to CryoEM or X-ray diffraction. Cationic, anionic, nonpolar and glycine residues are colored blue, red, black and green, respectively. **C.** ^1^HN-^1^Hα region of 2D ^1^H-^1^H COSY (maroon/dark green), TOCSY (blue) and NOESY (magenta) spectra. Inter-residue crosspeaks are colored maroon and intra-residues crosspeaks are colored black. **D.** ^1^HN-^1^Hα region of 2D ^1^H-^13^C HSQC spectrum.

**Supporting Figure 4:**
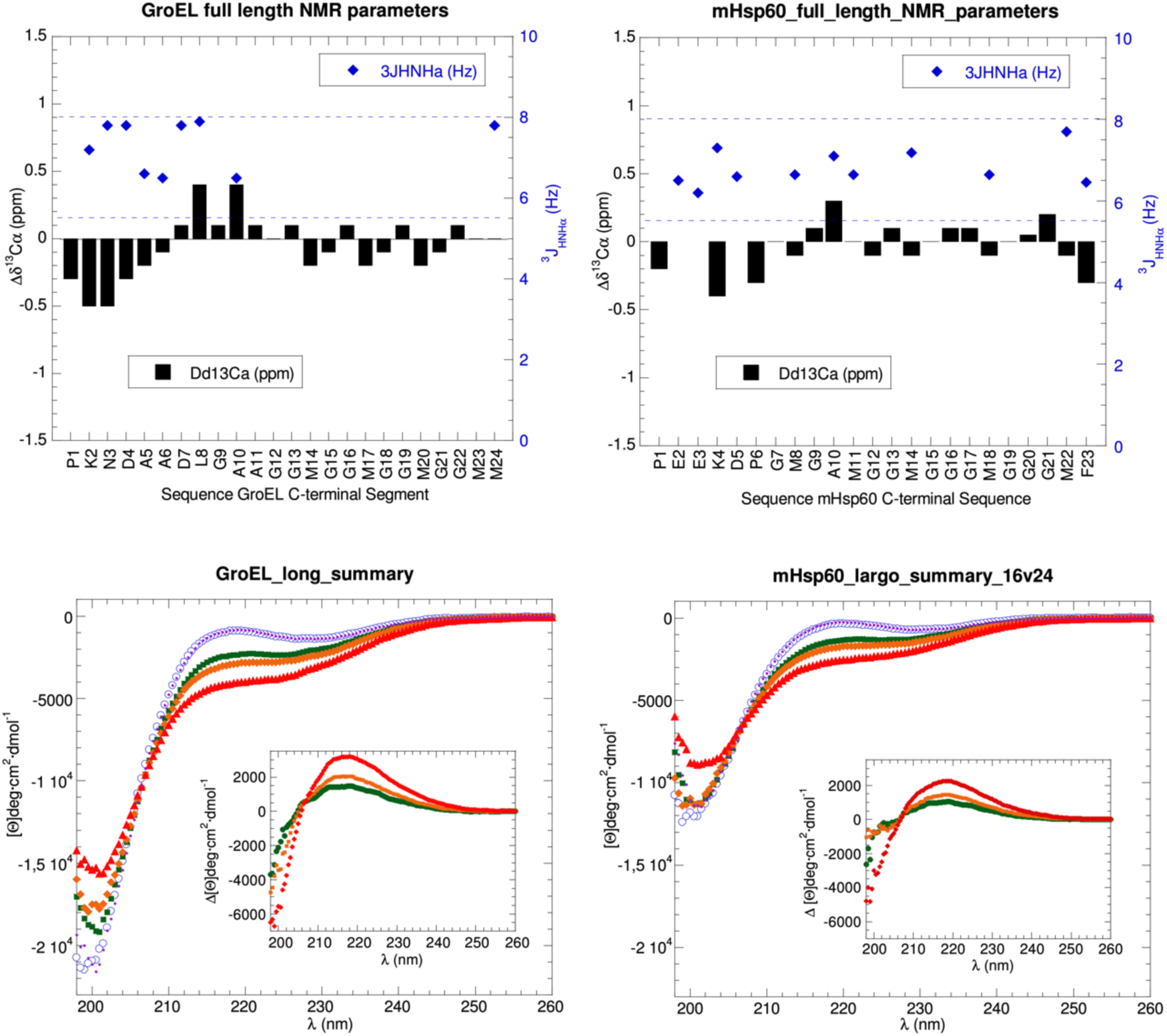
NMR Parameters and CD Spectra of GroEL and mHsp60 C-terminal Segment (complete) ^13^Cα conformational chemistry shifts (black bars, left y-axis) and ^3^J_HNHα_ coupling constants (blue diamonds, right y-axis) for the GroEL (top left panel) and mHsp60 (top right panel) C-terminal segments. Blue dashed horizontal lines mark the range of constants (between 5.5 & 8.0 Hz) expected for statistical coil or polyproline II conformations. Far UV-CD spectra recorded at 5°C (blue), 25°C (green) 37°C (orange) and 65°C (red) and after recooling to 5°C (purple) for the C-terminal tails of GroEL (bottom left panel) and mHsp60 (bottom right panel). Insets show the difference CD spectra for 5°C-25°C (green), 5°C-37°C (orange) and 5°C-65°C (red).

**Supporting Figure 5:**
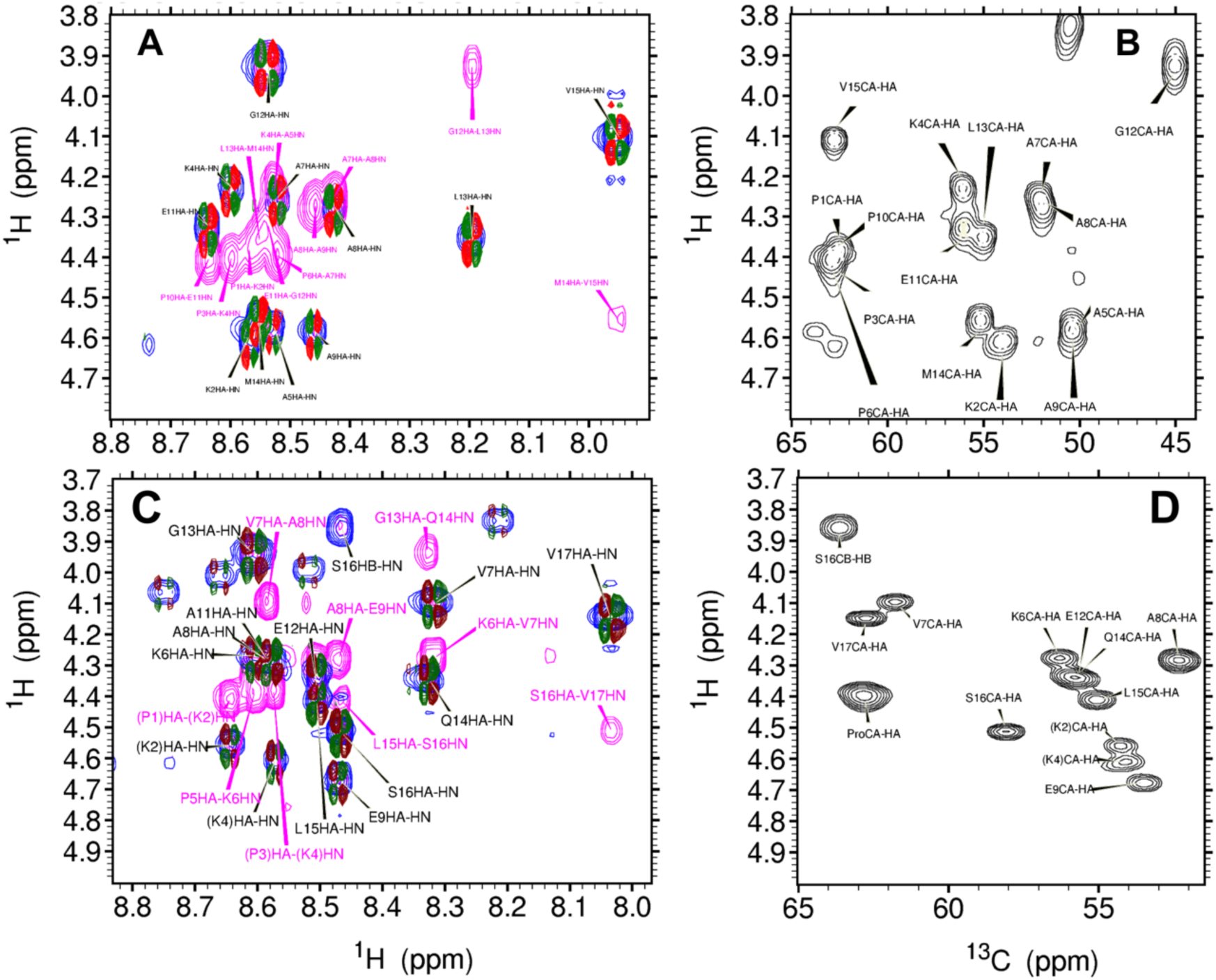
NMR Spectra of *Arabadopsis thaliana* and wheat Cpn60α C-terminal segments. **A.** ^1^HN-^1^Hα region of 2D ^1^H-^1^H TOCSY (blue), NOESY (magenta), and COSY (maroon/dark green) spectra of *A. thaliana* Cpn60α Ct. Intraresidue crosspeaks are labeled in black and sequential NOESY crosspeaks are labeled in magenta. **B.** ^1^Hα-^13^Cα region of the 2D ^1^H-^13^C HSQC spectrum of *A. thaliana* Cpn60α Ct. The signals arising from the proline residues are overlapped. Contours are plotted with a 1.4 multiplication factor. **C.** ^1^HN-^1^Hα region of 2D ^1^H-^1^H TOCSY (blue), NOESY (magenta), and COSY (maroon/dark green) spectra of Wheat Cpn60α Ct. Intraresidue crosspeaks are labeled in black and sequential NOESY crosspeaks are labeled in magenta. Assignments for K2 and K4 as well as P1 and P3 (in parentheses) are ambiguous. **D.** ^1^Hα-^13^Cα region of the 2D ^1^H-^13^C HSQC spectrum of Wheat Cpn60α Ct. Assignments for K2 and K4 (in parenthesis) are ambiguous and the signals arising from the proline residues (labeled “Pro”) are overlapped. Contours are plotted with a 1.4 multiplication factor.

**Supporting Figure 6.**
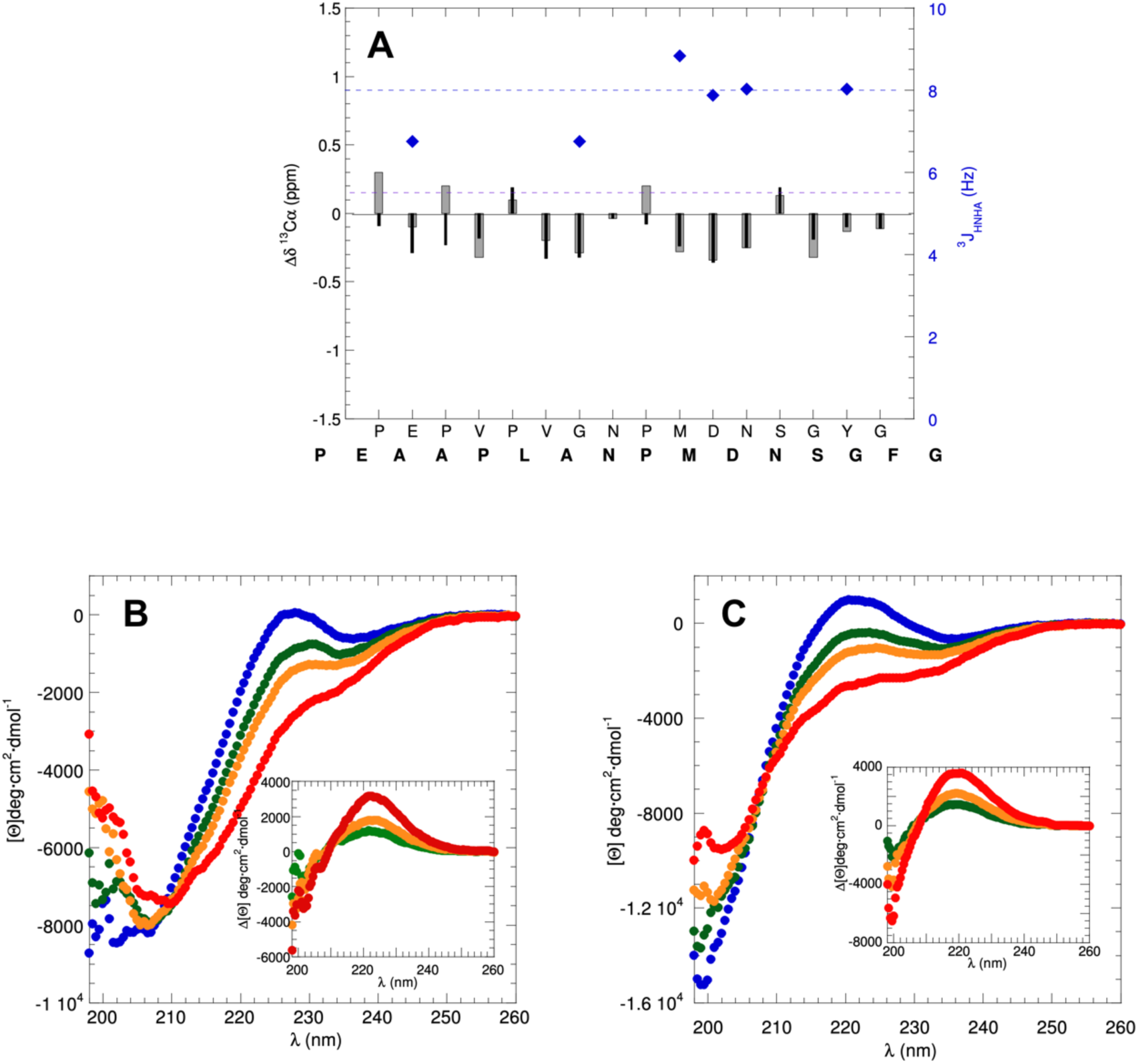
NMR Parameters and CD Spectra of *A. thaliana* and wheat Cpn60β. **A.** ^13^Cα conformational chemistry shifts of *A. thaliana* (black narrow bars, left y-axis) and wheat (gray wide bars, left y-axis) and wheat ^3^J_HNHα_ coupling constants (blue diamonds, right y-axis) Cpn60β C-terminal segments. Blue dashed horizontal lines mark the range of constants (between 5.5 & 8.0 Hz) expected for statistical coil or polyproline II conformations. Far UV-CD spectra recorded at 5°C (blue), 25°C (green) 37°C (orange) and 65°C (red) and after recooling to 5°C (purple) for the C-terminal tails of *A. thaliana* Cpn60β (panel **B**) and wheat Cpn60β (panel **C**). Insets show the difference CD spectra for 5°C-25°C (green), 5°C-37°C (orange) and 5°C-65°C (red).

